# Candidate gene length polymorphisms are linked to dispersive behaviour: searching for a mechanism behind the “paradox of the great speciators”

**DOI:** 10.1101/2023.01.19.524190

**Authors:** Andrea Estandía, Ashley T. Sendell-Price, Graeme Oatley, Fiona Robertson, Dominique Potvin, Melanie Massaro, Bruce C. Robertson, Sonya M. Clegg

## Abstract

The “paradox of the great speciators” has puzzled evolutionary biologists for over half a century. A great speciator requires excellent dispersal ability to explain its occurrence on multiple islands, but reduced dispersal ability to explain its high number of subspecies. A rapid reduction in dispersal ability is often invoked to solve this apparent paradox, but a proximate mechanism has not been identified. Here, we explore the role of six genes linked to migration and animal personality differences (*CREB1, CLOCK, ADCYAP1, NPAS2, DRD4*, and *SERT*) in 20 South Pacific populations of silvereye (*Zosterops lateralis*) that range from highly sedentary to partially migratory, to determine if genetic variation is associated with dispersal propensity. We detected genetic associations in three of the six genes: i) in a partial migrant population, migrant individuals had longer microsatellite alleles at the *CLOCK* gene compared to resident individuals from the same population; ii) *CREB1* displayed longer average microsatellite allele lengths in recently colonised island populations (< 200 years), compared to evolutionarily older populations. Bayesian broken stick regression models supported a reduction in *CREB1* length with time since colonisation and decreasing dispersal propensity; and iii) like *CREB1, DRD4* showed differences in polymorphisms between recent and old colonisations but a further sample size is needed to confirm. *ADCYAP1, SERT*, and *NPAS2* were variable but that variation was not associated with dispersal propensity. The association of genetic variants at three genes with migration and dispersal ability in silvereyes provides the impetus for further exploration of genetic mechanisms underlying dispersal shifts, and the prospect of resolving a long-running evolutionary paradox through a genetic lens.

## Introduction

Dispersal ability often determines the extent of species ranges and patterns of population divergence and speciation (Birand et al., 2012; Waters et al., 2020). Highly dispersive taxa tend to have larger geographic ranges (Lester et al., 2007; Sheard et al., 2020) and more opportunities to exchange genes among populations in different parts of their range, inhibiting diversification (Lack, 1969; Birand et al., 2012; Gillespie et al., 2012, but see Cardillo, 2003). In addition, the spatial scale of speciation is linked to variation in gene flow across a range of taxa (Kisel and Barraclough, 2010), and reduced gene flow during early stages of divergence is central to many speciation models (Hedrick, 1983; Slatkin, 1987; Coyne and Allen Orr, 1998; Mayr and Diamond, 2001; Grant and Grant, 2008; Price, 2008).

Great speciators are those bird species that inhabit multiple islands reached by overwater dispersal events, suggesting excellent colonisation ability, yet display multiple morphological subspecies (Diamond et al., 1976; Moyle et al., 2009; Pedersen et al., 2018; Manthey et al., 2020). This concept is applied to several subspecies-rich birds from Melanesia and the South Pacific, including representatives within the white-eyes, kingfishers, and cicadabirds (Mayr and Diamond, 2001). A common explanation to reconcile broad island distributions with phenotypically diverse taxa, is that dispersal traits are strongly selected against in each newly established island population (Diamond, 1981). Selection may reflect the individual energetic cost of this behaviour relative to the benefits of remaining (McNab, 2002; Bonte et al., 2012) or the idea that once most islands are filled, the chance of successful overwater dispersal is diminished (Diamond, 1970). However, it is still unclear how these relatively rapid, post-colonisation changes in dispersal propensity could arise.

Variation in dispersal propensity is largely determined by differences in a combination of morphological, physiological and behavioural traits (Matthysen, 2012). In birds, the types of morphological changes that indicate a reduced dispersal ability include a more rounded wing (as indicated by a lower hand-wing index) (Kipp, 1959; Lockwood et al., 1998) or a shift to a more graviportal body plan (reduced flight muscles and longer legs) (Wright et al., 2016). However, among the great speciators, the shift from dispersive to sedentary forms may primarily involve behavioural rather than morphological or physiological changes, at least initially (Diamond, 1981). This is termed “behavioural flightlessness”, a reluctance to disperse (especially across water), despite the maintenance of normal wings and flight. Hence, focusing solely on morphological proxies for dispersal ability that take time to change risks missing an essential part of the early evolutionary process.

A useful but relatively unexplored framework to understand behavioural flightlessness and subsequent divergence considers the links between dispersive behaviours and personality traits (Ingley and Johnson, 2014). Migration (seasonal movements) or dispersal (single movement to a new area) behaviours and personality traits are linked to some extent – migratory or highly dispersive taxa often show increased levels of boldness, aggression and exploration, and lower levels of sociability. Migratory individuals within a species tend to be bolder than non-migratory ones (Chapman et al., 2011). For example, highly aggressive Western bluebird (*Sialia mexicana*) individuals were found to be more dispersive than their less aggressive counterparts (Duckworth and Badyaev, 2007; Duckworth and Kruuk, 2009); European Stonechat (*Saxicola rubicola*) migrant males were more territorial than resident males (Marasco et al., 2011); and dispersive great tit (*Parus major*) individuals had a greater exploration rate than non-dispersive individuals (Korsten et al., 2013).

Both quantitative and molecular genetic studies provide evidence of a genetic influence on migratory and dispersive behaviours (Liedvogel et al., 2011). These behaviours often have significant narrow sense heritability, often greater than 0.4 (Dochtermann et al., 2019). Additionally, at the molecular level, genetic variation for migration, dispersive behaviour and personality traits are well documented (Bubac et al., 2020), including the identification of a number of candidate genes underlying personality traits such as tendency for boldness or explotatory behaviour (Fidler et al., 2007; Steinmeyer et al., 2009; Ruegg et al., 2014; Canestrelli et al., 2016). Changes in genetic variation at these candidate genes may act as a genetic switch catalysing population-level shifts in dispersive behaviour.

To date, six candidate genes are thought to contribute to a migratory phenotype, dispersive behaviour and personality traits in at least some bird species (Bubac et al., 2020; Table S1). In four of these genes, “adenylate cyclase activating polypeptide 1 gene” (*ADCYAP1);* the polyglutamine repeat region of the “circadian locomotor output cycles kaput gene” (*CLOCK;* Johnsen et al., 2007), the “neuronal PAS domain protein 2” (NPAS2), and the “cAMP responsive element binding protein 1” (CREB1)), microsatellite allele length variation is associated with migratory-related traits across a variety of avian taxa (Table S1). The other two genes (“dopamine receptor D4” (*DRD4*) and “serotonin transporter” (*SERT)*) show associations between single nucleotide polymorphisms (SNPs) and differences in avian personality and migratory propensity (Table S1).

The Zosteropidae family (white-eyes, yuhinas and allies) consists of 142 species (Clements et al., 2021) many of which are highly dispersive as evidenced by colonisation of numerous oceanic islands throughout the Indian and Pacific oceans, along with the broad continental distributions of some species (Mees, 1969; Clegg et al., 2002). This family shows one of the highest per-lineage diversification rates for vertebrates (Moyle et al., 2009) and divergence can occur even across minor geographic barriers (e.g. water gaps of just 2 km) (Mees, 1969; Moyle et al., 2009; Bertrand et al., 2014; Cowles and Uy, 2019; Manthey et al., 2020). A particularly interesting species within this family is the silvereye (*Zosterops lateralis*), considered a great speciator having multiple subspecies (17 morphological subspecies; Clements et al., 2021) with a wide natural distribution - including the Australian mainland and Tasmania, the North and South Islands of Aotearoa New Zealand (henceforth New Zealand), outlying oceanic islands of Australia and New Zealand, and the archipelagos of New Caledonia, Vanuatu, and Fiji (Fig. 1A). Silvereyes also display a variety of gene flow potentials: the Tasmanian subspecies (*Z. l. lateralis*) is a partial migrant (Mees, 1969); those in the central Vanuatu archipelago display high levels of outgoing gene flow, while more peripheral populations have high levels of incoming gene flow (Clegg and Phillimore, 2010); and others, such as on Heron Island and Lord Howe Island in the southern Great Barrier Reef, are sedentary and genetically isolated (Sendell-Price et al., 2020).

**Fig. 1.**
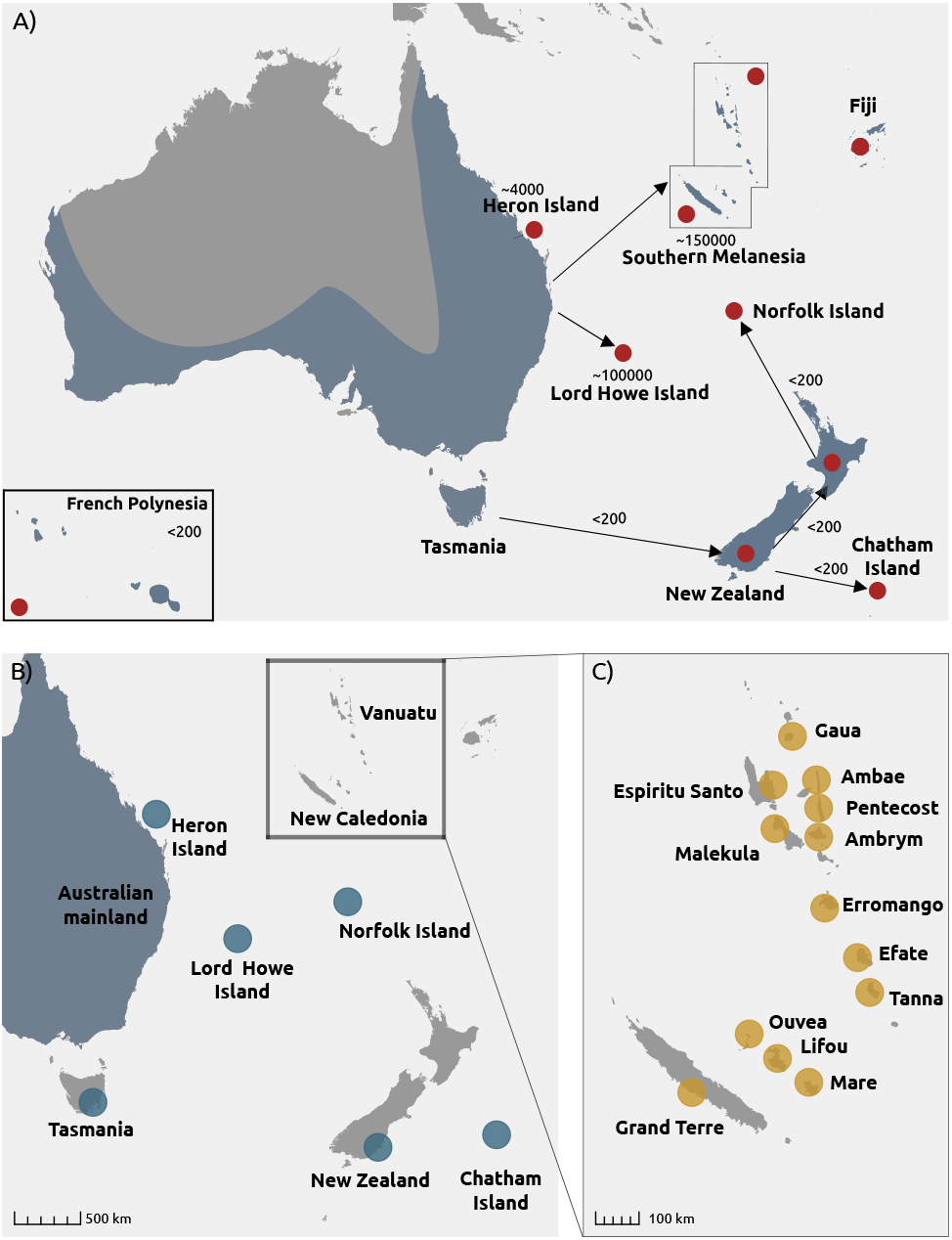
A) Distribution range of the silvereye highlighted in blue. Red dots represent single islands or archipelagos where multiple islands have been colonised from the Australian mainland or Tasmania. Approximate colonisation times are shown. French Polynesia is shown as inset; Locations of silvereye populations sampled for candidate gene variation: B) ANZO cluster C) SM populations.

Here, we assess variation in six personality-related candidate genes in the silvereye examining different levels: individuals, populations and subspecies. Our aim is to determine whether there are signatures consistent with a genetic switch that would explain rapid shifts in dispersal, leading to reduced gene flow and subsequent increased opportunity for divergence. We ask i) if candidate gene variation follows patterns of neutral genomic divergence, that reflect drift and gene flow processes only; ii) if candidate gene variation is correlated with dispersal propensity among a range of dispersive and non-dispersive populations of silvereye, considering time since colonisation; and iii) if candidate gene variation is correlated with individual migratory status in the partially migratory Tasmanian population.

## Methods

### Sampling and DNA Extraction

Silvereye blood samples were collected between 1996 and 2016 from 20 sites in eastern Australia and New Zealand, including outlying islands, henceforth ANZO (Fig. 1B), and Southern Melanesia (New Caledonia and Vanuatu) referred to henceforth as SM (Fig. 1C; Table S3). The sampling included putative Tasmanian winter migrant individuals captured in the Australian mainland states of Queensland and New South Wales. Individuals were assigned as Tasmanian migrants based on plumage differences: resident silvereyes (subspecies Z. *l. cornwalli*) exhibit bright yellow throat and grey flanks, while migrant individuals (subspecies Z. l. *lateralis*) exhibit white-to-pale-yellow throat and dark buff flanks (Fig. S1; Higgins et al., 2006). Birds were caught in mistnets or traps and blood samples were taken via venipuncture of the brachial wing vein and stored in 90% ethanol or lysis buffer (10mM EDTA pH 8.0, 10mM TRIS-HCl pH 8.0, 20mM NaCl, 1% SDS, Seutin et al., 1991).

### Candidate microsatellite genotyping

We extracted DNA from 422 samples using a standard chelex protocol (Walsh et al., 1991) and amplified four microsatellite candidate genes (*NPAS2, CREB1, CLOCK*, and *ADCYAP1*) (Table S3) with the primers from Steinmeyer et al., (2009) using a touchdown polymerase chain reaction (PCR) protocol and fluorescent labelling (VIC and FAM) using M13 tags (Schuelke, 2000). Fluorescent labelling with VIC and FAM was done in multiplex PCR reactions containing two loci each (VIC: *CLOCK* and *ADCYAP1*; FAM: *NPAS2* and *CREB1);* final reaction volume of 3 μL contained 10ng DNA, 1 μl Type-it Master Mix (Qiagen), 0.17 μM of either FAM or VIC, and 0.01 μM forward M13-labelled primer and 0.05 μM reverse primer for each locus.

Thermal cycling consisted of an initial denaturation step of 95°C for 15 min, eight cycles of 94°C for 30s, an annealing temperature of 60°C, reduced by 1°C each cycle, for 90s and a 72°C extension step for 60s, then 25 cycles of 94°C for 30s, 52°C for 90s, and a 72°C for 60s, and a final extension of 60°C for 30 min. VIC and FAM multiplexes were pooled for each sample and allele sizes for the candidate gene microsatellites were determined in relation to LIZ600 size standard on an ABI 3730xl DNA Analyser and scored using the Microsatellite Plugin within Geneious 2020.1 (http://www.geneious.com, Kearse et al., 2012). We tested for the deviations from Hardy-Weinberg Equilibrium, allelic richness and heterozygosity using the R package *diverRsity* (Keenan et al., 2013).

### *DRD4* and *SERT* sequencing

We targeted one region of *DRD4*, encompassing intron 2 and exon 3, and part of intron 3 (1480 bp). For *SERT*, we amplified part of the promoter region (508 bp). Each 25 μl PCR reaction was comprised of approximately 10ng DNA template, 1.0 unit BioTaq (Bioline USA Inc.), 1 x BioTaq reaction buffer, and 0.5μM each forward and reverse primer, 200μM dNTPs, 1.5mM MgCl2, and made up to volume with Milli Q water. The PCR reaction profiles for both *DRD4* and *SERT* fragments consisted of an initial denaturation step of 94°C for 3 min, followed by 10 cycles of 94°C for 3 min, a touchdown step at 65°C for 30s, and an extension step of 72°C for 2 min. This was followed by 25 cycles of 94°C for 30s, 55°C annealing step for 30s, an extension step of 72°C for 2 min, and a final extension step of 72°C for 10 min. We purified PCR products using Acroprep 96 filter plates (Pall Corporation) following the manufacturer’s protocol, then Sanger sequenced with forward and reverse primers on an ABI 3730xl DNA Analyser (Genetics Analysis Service, Otago University, NZ). We aligned the sequences using MUSCLE (Edgar, 2004) in Geneious version 2020.1, and identified SNPs and insertions/deletions (indels). For *DRD4*, we aligned sequences to the great tit *DRD4* gene sequence (Genbank accession no.: DQ006801.1; Fidler et al., 2007) and *SERT*, we used the dunnock sequence (Genbank accession no.: KT967954.1; Holtmann et al., 2016). We called SNPs with a minimum minor allele frequency (MAF) of 0.05.

### Gene flow as a proxy for dispersiveness

As a proxy of population dispersiveness, we estimated outgoing gene flow rates from each population. We used a subset of whole genome sequences available from Estandía et al. (in prep) to examine population structure patterns in NGSadmix (Skotte et al., 2013), a module implemented in ANGSD (Korneliussen et al., 2014). These whole genome sequences covered 336 of 422 individuals included in the present study. We generated a BEAGLE file containing genotype likelihoods and created a subset of 10,000 SNPs picked at random after applying filtering for a MAF of 0.05. We ran NGSadmix with a range of genetic clusters (*k*), from 2 to 20. We selected the best k for our dataset based on the mean estimate likelihoods, which indicated *k*=2 as the optimal number of clusters, corresponding to ANZO and SM groupings. We reran NGSAdmix in each of the main cluster to explore potential substructuring. We visualised clustering patterns using a custom R script.

We generated a covariance matrix in PCAngsd (Meisner and Albrechtsen, 2018) using the BEAGLE file. Because not all individuals were screened for all candidate genes (e.g. 375 individuals for *NPAS2* but 258 for *CLOCK*), and not all individuals in the candidate gene and whole-genome dataset coincide, we produced a population-level covariance matrix. From this, we calculated contemporary rates of gene flow among *Z. lateralis* populations for each of the two clusters (ANZO and SM) separately using BA3-SNP (Mussmann et al., 2019), an extension of BayesAss that allows SNPs as input. As input, we used the same file as for the population structure analysis (10,000 independent SNPs). This software employs a Bayesian approach with Markov chain Monte-Carlo (MCMC) sampling to estimate migration rates. We adjusted delta values for migration rates (m), allele frequencies (a) and inbreeding coefficients (f) to ensure that parameter space sampling acceptance rate was between 20% and 60% (Wilson and Rannala, 2003). We ran the program for one million iterations, discarding the first 10% as burnin. We estimated the 95% credible sets by calculating the mean±1.96*Standard deviation (SD).

To assess model convergence, we compared results from 10 replicate runs each with a different random starting seed. We considered runs to have converged on a similar solution if gene flow estimates were within 0.005 (0.5%) across runs. Additionally, we inspected the likelihood trace files in Tracer v 1.7 (Rambaut et al., 2018) to determine that the log of the posterior probability values were consistent and to ensure that the Effective Sample Size (ESS) values were greater than 200. The run with the highest log posterior probability value was considered the best run to obtain parameter estimates.

### Candidate gene association analysis

For all candidate genes we assessed variation among populations using the R package *brms* (Bürkner, 2017). Individual mean microsatellite length was set as the response variable and population was set as a categorical explanatory variable. In the case of *SERT* and *DRD4*, the response variable was coded as 0 or 1 representing whether the individual carried the most frequent nucleotide or the variant, and we used a bernoulli family for binary outcomes. For the two microsatellite candidate genes that showed obvious within and between population variation in length (*ADCYAP1* and *CREB1*) we tested whether population age and dispersal propensity could explain this variation by running Bayesian linear mixed models in *brms. CREB1* showed a clear distinction in allele lengths between a grouping consisting of the Australian mainland, Tasmania and recently colonised islands versus old island populations. Because of this structure (that does not completely align with ANZO and SM neutral structure groupings), we also applied a broken stick regression model for *CREB1* only to test whether including a single change point would improve predictive performance over an intercept-only and a linear model using *mcp* (multiple point change) (Lindeløv, 2020). *NPAS2* and *CLOCK* displayed little among population variation in average allele length, hence we did not apply Bayesian linear mixed models to the whole population set. However, *CLOCK* showed variation within the partial migrant population of Tasmania, so we tested whether individuals that migrated to the mainland had different *CLOCK* lengths to those that over-wintered in Tasmania. The Tasmanian sample was restricted to 14 winter-caught birds (non-migrants) as the resident summer population includes a mix of migrants and non-migrants that cannot be phenotypically distinguished. The ‘migrant’ group consisted of those caught at Australian mainland sites in winter that were phenotypically identified as Tasmanian migrants (26 individuals).

For Bayesian linear mixed models (*CREB1* and *ADCYAP1*) and broken-stick regression models (*CREB1*), the following population-level (fixed) parameters used were: (a) Dispersal Index (*DI*): the sum of each outgoing gene flow estimate (g) from island i into island j (where 95% credible interval did not overlap with zero), multiplied by the geographic distance between the islands (d) (Equation 1). The latter helps to account for differences in geographic opportunity for dispersal; for example, a geographically isolated island population that has moderate outgoing gene flow to few far islands would score higher than a centrally located island population with moderate outgoing gene flow to many close islands. *DI* was scaled from 0 (non-dispersive) to 1 (maximally dispersive).

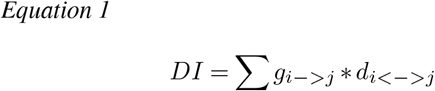

(b) Age (Table S2): population ages for New Zealand, Chatham Island and Norfolk Island are known from historical records (c. 190 years; Clegg et al., 2002; Mees, 1969); for Heron Island, the population age was set as a maximum of 4000 years based on length of time the island has been vegetated (Clegg et al., 2008); and molecular estimates for the remaining ancient populations were taken from a time-calibrated gene tree (Black, 2010).

The intercept-only model represents the starting hypothesis that mean *CREB1* or *ADCYAP1* length differs between populations (which was set as group-level parameter or random effect) but does not change as a function of increasing dispersal propensity or time since colonisation. We controlled for population structure using the PCAngsd population-level covariance matrix. As *mcp* does not permit incorporation of distance matrices, we used a categorical variable that classified each individual according to its membership in one of the two population genetic clusters (ANZO or SM) identified from the top-level NGSadmix analysis.

For all *brms* (a) Bayesian linear mixed model: Differences in length among populations for each candidate microsatellites; b) Bayesian linear mixed models: i) migration: *CLOCK*, and ii) population age and DI: *CREB1* and *ADCYAP1*) and *mcp* (*CREB1* only)), average lengths were modelled with a Gaussian distribution and application of the default link function. We set weakly informative priors for the intercepts of the segments before and after the change point (*mcp*). The same prior was applied for both *CREB1* length intercepts due to the lack of a priori information about a positive or negative correlation of mean *CREB1* length with population age and dispersal index. Priors for both intercepts were centred at 550 bp with a SD of 20 (N(550, 20)), based on known mean and variance of *CREB1* lengths in other passerines (Steinmeyer et al., 2009). For the *brms* linear mixed model, we set weakly informative priors with a normal distribution centred on 0 with a standard deviation of 20 for the DI and population age coefficients. This prior does not assume an increase or decrease in length, but incorporates the prior information that changes will not be greater than 20 bp. We performed prior predictive checks, where data was generated according to the specified prior predictive distributions in order to assess their suitability (Gelman et al., 2019).

For both the *brms* and *mcp* analyses we used MCMC with four chains of 4000 iterations each, including a warm-up of 400 iterations. We evaluated convergence via visual inspection of the MCMC trace plots, checking that the ESS>200 and the R values for each parameter (R = 1 at convergence). To evaluate model performance, we compared our fitted models with an intercept-only model using leave-one-out crossvalidation (LOO), a robust, fully Bayesian model selection approach (Vehtari et al., 2017).

## Results

### Population genetic structure and gene flow

NGSadmix analysis of WGS data supported two main genetic clusters (*k*=2): Cluster 1 comprised Australia, New Zealand and outlying island populations (ANZO), and Cluster 2 comprised Vanuatu and New Caledonia populations in Southern Melanesia (SM) (Fig. 2; Fig. S2; Table S4). Other values of *k* also had high likelihoods (Table S4); *k*=3 indicated substructuring within ANZO due to separation of Heron Island and Lord Howe Island from other populations (Fig. S2; see also *k*=2 in the ANZO analysis Table S4.1); *k*=4 indicated sub-structuring within the SM cluster, primarily separating New Caledonia from Vanuatu populations, with the Vanuatu island of Tanna showing some affiliation with New Caledonia (Fig. S2, see also *k*=2 in the SM analysis Table S4.2). These population genetic patterns were consistent with those that emerged from the covariance matrix (Table S5, Fig. S3). The ten independent BayesAss runs conducted to quantify the degree and direction of migration rates all agreed on the patterns of gene flow (Table S6). Of 42 pairwise comparisons within the ANZO cluster, and 132 in the SM cluster, eight and six respectively had estimates for which the credibility intervals did not overlap with zero. Within ANZO, this was primarily seen in relatively high outgoing gene flow estimates from Tasmania and New Zealand, and within SM, moderate outgoing levels from central islands of Pentecost and Malekula.

**Fig. 2.**
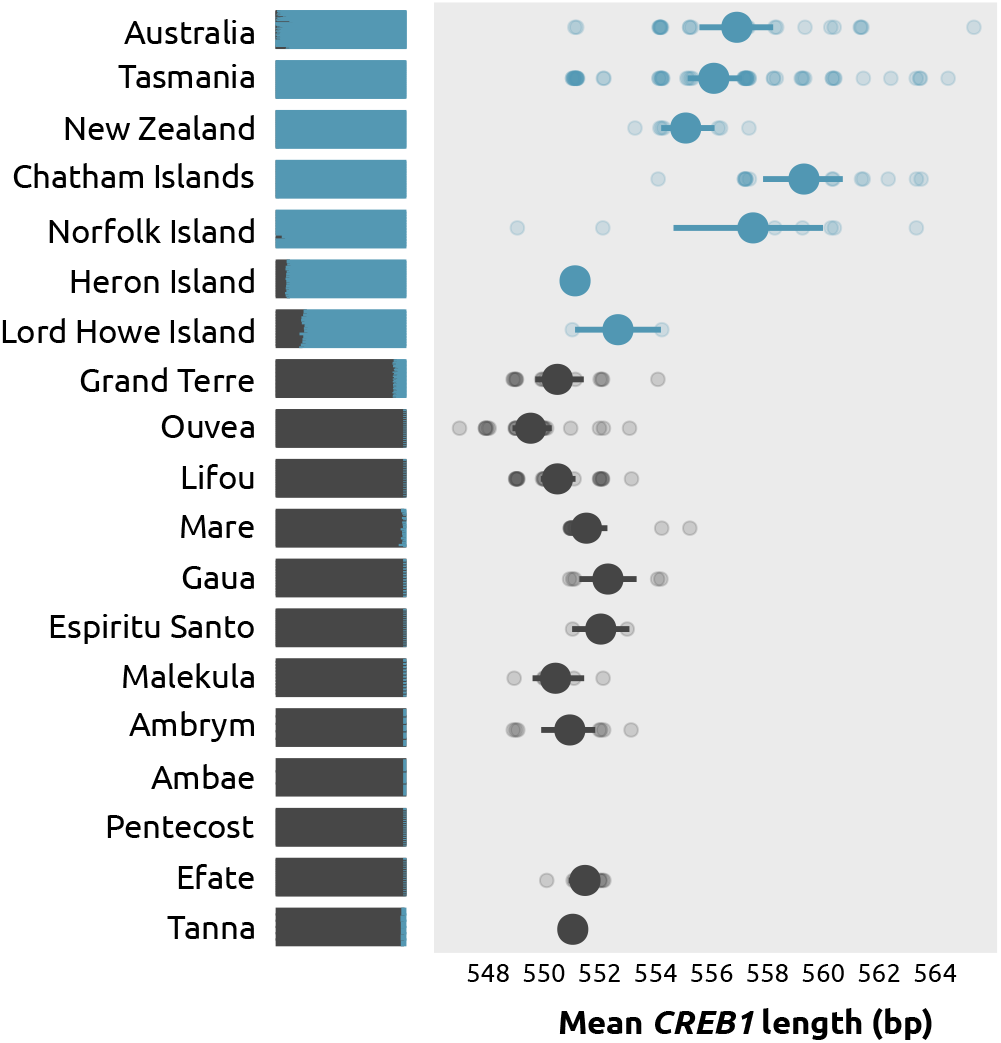
A) NGSadmix plot (k = 2) showing a split between ANZO and the SM populations. B) The average *CREB1* allele length per population indicates a reduction in length for Heron and Lord Howe Islands, and all populations in the SM cluster. *CREB1* was not screened for Ambae and Pentecost samples.

### Variation in candidate microsatellites, *DRD4* and *SERT*

Mean and individual allele lengths for microsatellite candidate loci are shown in Figure 2 and Figure 3. *CREB1* showed longer mean lengths for the Australian mainland, Tasmania (migrants and non-migrants), and the recently colonised populations of New Zealand, Chatham Island, and Norfolk Island (Fig. S4; Table S7.1). Heron Island, Lord Howe Island and all Southern Melanesian populations displayed shorter allele lengths on average.

**Fig. 3.**
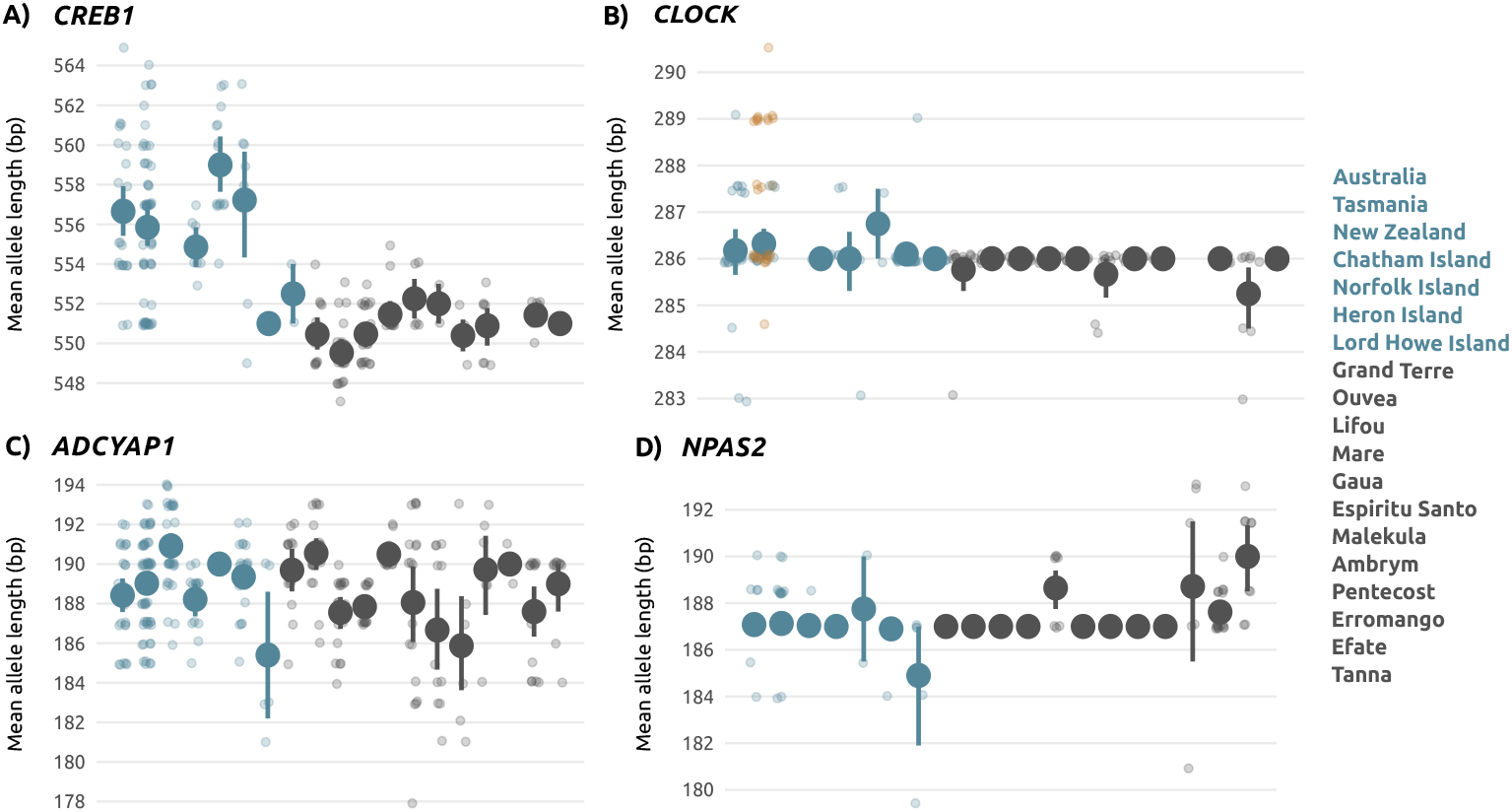
Mean individual allele lengths (shadowed dots) with mean and standard error for each population (large dots) for the four microsatellite candidate genes. Blue and grey dots correspond to two population genetic clusters identified from admixture analysis. A) *CREB1* is longer in the source populations (Australia and Tasmania), and recently colonised populations (New Zealand, Chatham Islands and Norfolk Island) compared to older populations (Heron Island, Lord Howe Island, and SM populations). B) Migrant individuals highlighted in orange have longer allele lengths at *CLOCK*. C) *ADCYAP1* shows extensive variation, including shorter lengths in Lord Howe Island and Malekula and longer lengths in New Zealand, Ouvea and Gaua. D)*NPAS2* shows little variation with southern Vanuatu populations (Erromango, Efate and Tanna) showing an increased mean allele length.

*CLOCK* was monotypic in the majority of populations (Fig. 3B; Table S7.2). Compared to Tasmanian resident silvereyes, Tasmanian migrants showed longer allele lengths. Migrant individuals had long *CLOCK* variants (allele lengths of 289 and 291) not observed in any winter-caught Tasmanian birds (i.e. residents) (Fig. 4A), or any other population except for a Heron Island individual. *NPAS2* showed some variation across populations but similar mean values for populations in the ANZO cluster (Fig. 3D; Table S7.3). However, most SM populations were not variable at this locus, with the exception of peripherally located islands of Gaua, Efate and Tanna in Vanuatu. Variation between and within populations for *ADCYAP1* was evident for all populations (Fig. 3C; Table S7.4).

**Fig. 4.**
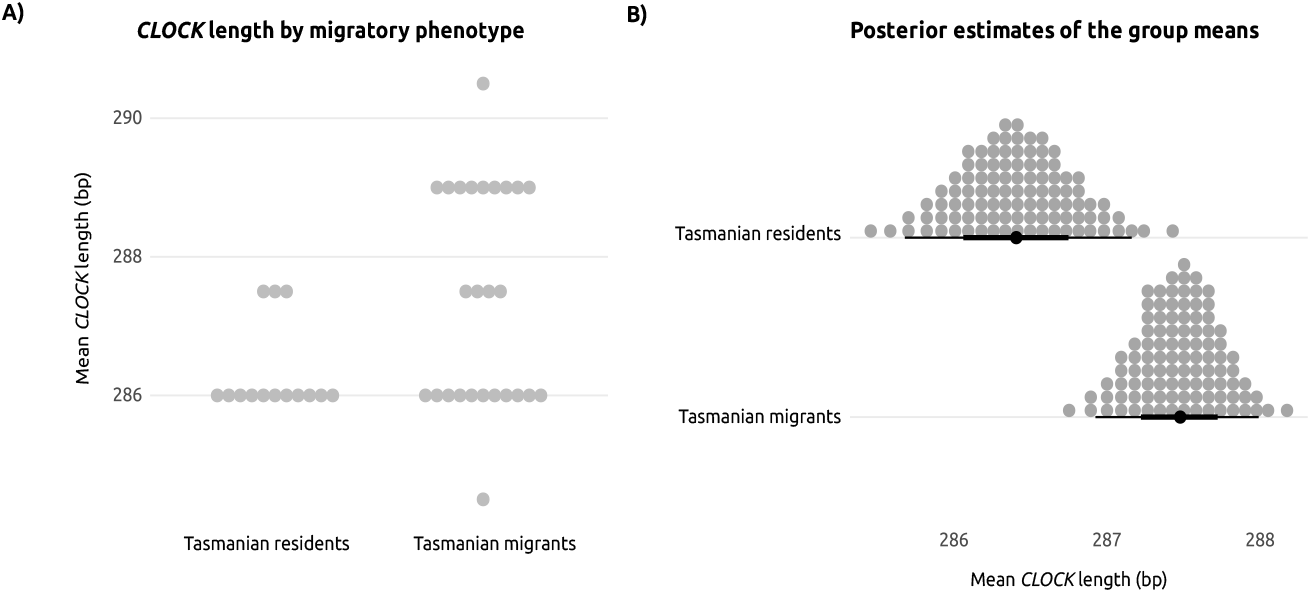
A) Individual *CLOCK* lengths for Tasmanian residents and Tasmanian migrants (Tasmanian silvereyes caught in winter on the Australian mainland). Tasmanian migrants show longer allele lengths. B) Posterior estimates of the group means obtained from the *brms* model. Migrant individuals have a 1bp increase when compared with residents.

We identified ten *DRD4* SNPs, seven of which were non-synonymous and three synonymous. Five of them showed significant differences across populations but only SNP at position 83 (a non-synonymous substitution) displayed consistent differences between ANZO and SM clusters (Table S7.5). SM individuals exclusively carried adenines, resulting in production of lysines, while those from New Zealand and Chatham Island only carried guanines producing argines. Tasmanian residents and Tasmanian migrants had both nucleotides represented. We found one SNP and an INDEL in *SERT* but none were significantly different in frequency among populations.

### Candidate gene association tests

Tasmanian migrants showed an increase of 1bp relative to non-migrants (Fig. 4B; Table S8). Mean *CREB1* variation was better explained by a single change point model than the intercept-only or a linear regression model (Fig. S4; Tables S9.1-S9.2). Time since colonisation and *DI* failed to explain the variability in *ADCYAP1* (Tables S10.1-S10.2).

The single change point model indicated that mean *CREB1* length decreased six base pairs with increasing population age, however the timing of this change had high uncertainty (Intercept 1 = 552.462, Intercept 2 = 557.494; Table S10.1; Fig. 5A). The posterior probability density of change point ranged between 200 and 4000 years ago when we see the step reduction in allele length in the Heron Island population (maximum age 4000 years, Clegg et al. 2008), a reduction that is observed in other older populations (Lord Howe Island and Southern Melanesian populations). The exception is Tasmania, an evolutionarily old population with long average *CREB1* length. *CREB1* decreased five base pairs with decreasing *DI* (Intercept 1 = 551.22, Intercept 2 = 556.68; Table S10.2; Fig. 5B). The posterior probability density of change point for *DI* ranged from Chatham Island to Efate, an island in Vanuatu that shows low levels of outgoing gene flow into nearby islands.

**Fig. 5.**
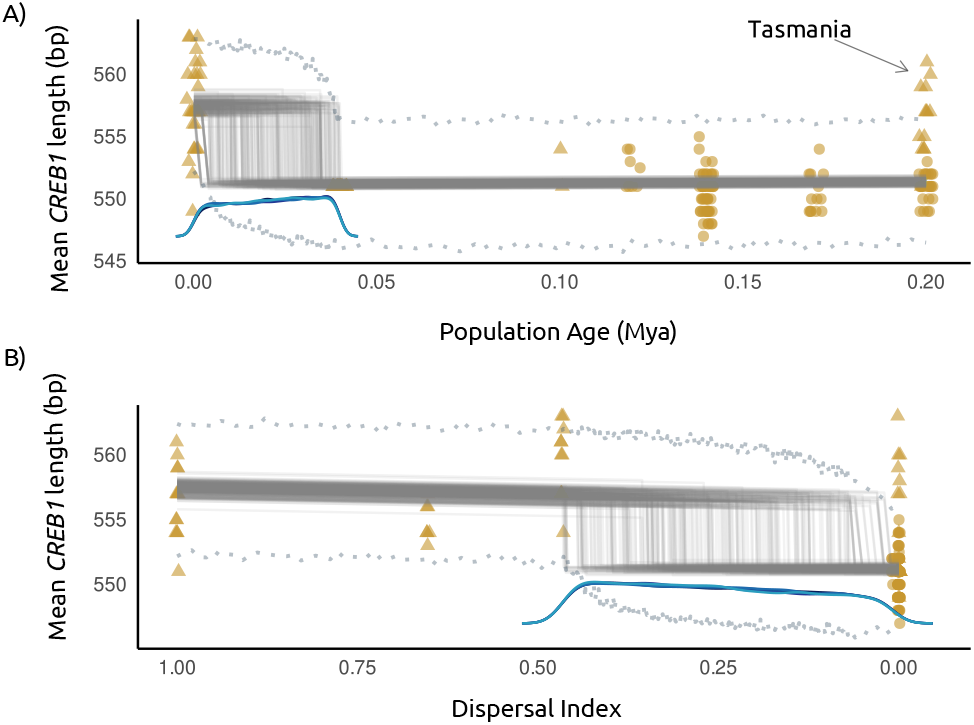
Broken-stick regression model for *CREB1* allele length variation. ANZO population data points (triangles) and SM population data points (circles); 100 posterior draws (grey lines); change point posterior distributions (four blue density curves, each representing one chain); and 89% prediction intervals (grey dashed lines). A) Relationship with population age, showing a higher mean length in recently colonised populations and their Tasmanian source population. Change point posteriors range from 200 years ago to 4000 years ago. B) Relationship with dispersal index (DI). The highest DI corresponds to Tasmania followed by New Zealand and Chatham Islands. Norfolk Island is among the young populations with no outgoing gene flow but long mean *CREB1* length. The change point posterior is centred between the DI values for Chatham Islands and Efate.

## Discussion

Rapid changes in dispersal could help to explain why taxa present on many islands show high levels of phenotypic diversity – the paradox of the great speciators. Quantifying variation in six candidate genes thought to underlie dispersal and migratory behaviour in populations of Australian mainland and island-colonising silvereyes has revealed three key results that hint at a role for genetic changes as a mechanism underlying this paradox. First, at the population level, we found that *CREB1* allele length variation was associated with time since colonisation and dispersal propensity, with recently colonised populations and those with high dispersal tendencies carrying longer alleles. Second, at the individual level, we found allele length variation in *CLOCK* associated with migratory status when comparing Tasmanian residents and Tasmanian migrants. Third, more limited data from a single *DRD4* SNP was fixed for different alleles in recently colonised versus evolutionarily older island populations. Together, these genes are likely to provide useful signatures of behavioural shifts in dispersal propensity in other silvereye populations, and possibly closely related species, though whether any of these genes act as a genetic switch remains unclear.

In the silvereye populations studied here, *CREB1* showed a disjunct pattern of allele sizes: longer in the mainland, the partial migrant Tasmanian population, and all of the recently colonised populations and shorter in island populations colonised thousands, or hundreds of thousands of years ago. This cannot be explained by population genetic groupings, as both the Heron Island population (up to 4000 years old) and Lord Howe Island population (over 100 thousand years old) (Clegg et al., 2002; Sendell-Price et al., 2020) had shorter *CREB1* allele lengths, yet clearly fell in the ANZO population genetic cluster that includes all of the long-allele-length populations. The Heron Island samples were monomorphic for a unique (551bp) allele, and we cannot rule out that it may be fixed entirely due drift in this small population (of approximately 300 breeding birds, McCallum et al., 2000). However, at the broad scale, the most dispersive populations, and those evolutionarily recent colonisation events, carry longer allele lengths, compared to more sedentary, and more ancient island populations. This is the opposite pattern to that found in common buzzard (*Buteo buteo*) where non-dispersive individuals carried longer *CREB1* allele lengths (Chakarov et al., 2013). *CREB1* allele lengths have also been shown to correlate with other life-history traits that also depend on circadian and circannual clocks e.g. incubation duration (Bourret and Garant, 2015) and male moult speed (Bazzi et al., 2017). The broad scale patterns, and suggested association with dispersal, were not reflected within each of the longer- and shorter-allele length groups. For instance, in the short allele length group, Gaua and Espiritu Santo displayed the longest lengths, but according to our gene flow analyses, they are not the most dispersive populations within their groupings; likewise for Norfolk Island in the long allele length group.

While the obvious length variation pattern seen at *CREB1* appears to have biological relevance for dispersal propensity in our study system, a mechanistic resolution of the paradox of the great speciators also requires dispersal shifts to be rapid (Diamond et al. 1976). We were unable to pinpoint when the *CREB1* allele size reduction occurs because of a lack of island populations of intermediate ages. At best we can say that for silvereyes, a shift in dispersal propensity is relatively rapid, taking more than 63 to 95 generations (South Island New Zealand, 190 years since colonisation, generation time of 2 to 3 years) and less than one to two thousand generations (Heron Island, 3,000 to 4,000 years old) (Clegg et al., 2008). Variation at a second gene, *CLOCK*, was associated with migratory status of individuals, with migrant individuals having longer *CLOCK* allele variants, including long variants absent in individuals from non-migrant individuals from Tasmania, Australian mainland and all island populations. Previous studies in migratory birds focusing on *CLOCK* variation show mixed results. Longer allele lengths have been found to be positively correlated with migratory propensity (Peterson et al., 2013), and variation in the phenology of migratory species (Liedvogel et al., 2009; Caprioli et al., 2012; Bazzi et al., 2015), but in some cases allele lengths are negatively correlated with migration date (Ralston et al., 2019) or not correlated at all (Mueller et al., 2011; Contina et al., 2018; Parody-Merino et al., 2019). *CLOCK* plays a key role in regulating the circadian oscillator gene complex (Panda et al., 2002), and is associated with variation in the phenology of photoperiodic traits (e.g. migratory behaviour) (Table S1). Photoperiodic stimulation experiments on caged Tasmanian silvereyes resulted in migratory restlessness being triggered only in migratory caged birds but not in resident ones, supporting a genetic link with migratory behaviour (Chan, 1994). Consequently, photoperiod changes could be the dispersal trigger with the onset of shorter autumnal days. The Tasmanian population is one of the few silvereye migrant populations situated where changes in day length during the nonbreeding, winter period is substantially greater than changes experienced further north and closer to the equator. Another location where these conditions are met is New Zealand. In fact, even though records are few, large flocks of silvereyes seem to have migrated from the South to the North Island (Dennison et al., 1981), providing further support that photoperiod changes which are mediated at least partially by genetics, can lead to migration. Some overwater dispersal events might be a direct consequence of off-course migration.

For example, the recent sequential colonisation sequence of the silvereye from Tasmania to New Zealand and outlying islands was likely initiated by an off-course flock of Tasmanian migrants (Mees 1969). The Tasmanian migrants sampled in this study carry long *CLOCK* allele variants that were not recovered in the recently colonised populations. This could be a consequence of our limited sample size, and/or underlying effects of a different set of genes that we have not included in this study on dispersal ability in these populations. The genetic basis of partial migration in Tasmanian silvereyes and whether associated variants are present in recently colonised populations should be further explored in future work using a hypothesis-free genome-wide approach.

*DRD4* is one of the most well-studied candidate genes related to exploratory and risk-taking behaviour (Bubac et al. 2020) – both of which could feasibly have links to dispersal propensity. We found a single polymorphism within *DRD4* (SNP83) that showed fixed differences between two young (more dispersive) and seven old (more sedentary) island populations. The Tasmanian population was polymorphic (G and A represented), the recently established populations of Chatham Island and South Island New Zealand were fixed for A, and Southern Melanesian populations fixed for G. SNP83 is located in intron 2 and corresponds to base pair 9,423 on the *DRD4* orthologue of the great tit (Fidler et al., 2007). It has not been noted previously as having any phenotypic associations. Previous work assessing *DRD4* variation and personality in the great tit focused on associations between exploratory behaviour and variation at ‘SNP830’, revealing large effects in certain populations but not in others (Fidler et al., 2007; Korsten et al., 2013; Riyahi et al., 2017), however SNP830 was not variable in our dataset. Our data adds to the evidence that variability in the *DRD4* gene plays a role in a suite of behavioural phenotypes, however the extent of importance of SNP83 will require screening more individuals and populations in silvereyes and other species.

Even though *NPAS2*, *ACDYAP1* and *SERT* each showed some variability between populations this variability not related to dispersal ability. Given the limitations of studying a handful of candidate genes for explaining complex behavioural phenotypes, failure to detect associations is not entirely unexpected. In birds, different associations (negative, positive or no correlation) in different populations, species and candidate genes are often reported (Table S1). Furthermore, few studies have considered candidate gene-environment interactions (but see Liedvogel et al., 2009; Liedvogel and Sheldon, 2010; Bourret and Garant, 2015) or methylation patterns that have been found to explain diverse complex behaviours in birds (Saino et al., 2019). The lack of replicability even within the same species could also be a product of sampling in different locations with a low number of individuals leading to a lack of statistical power, choosing different variables and proxies to measure migration and dispersiveness, using different methodologies to analyse data, publication bias or a combination of all of these (Yang et al., 2022). Thus, the lack of association in *NPAS2*, *ADCYAP1* and *SERT* does not rule out their potential role in dispersal behaviour as other factors might be masking their effects.

### Maintenance of high dispersal propensity in a continental island population

Despite being an old insular form (>200K split from Australian mainland subspecies (Black, 2010)), the Tasmanian silvereye (*Z. l. lateralis*) has maintained high dispersal propensity: it is a partial winter migrant to mainland Australia, it was the original source population for the historical sequential colonisation of New Zealand and outlying islands (Clegg et al., 2002; Mees, 1969), and as shown here, displays high levels of gene flow with Australian mainland subspecies. It also maintains the longer *CREB1* average allele lengths, and putative Tasmanian migrants caught on the mainland have shown unusually long *CLOCK* mean allele lengths. The maintenance of high dispersal potential, and its partial migrant status, are most likely explained by its geography and history of connectivity with the mainland. Tasmania has repeatedly been connected to the Australian mainland during glacial periods and it is currently separated by a very shallow sea (average depth: 60 m and shortest distance to the Australian mainland: 200 km; Blom and Alsop, 1988). Over 50 islands of varying sizes can be found between the Australian mainland and Tasmania, which can act as migration stopovers, and facilitate connectivity (Belbin et al., 2021).

### The future of dispersal genomics

A candidate gene approach to understanding the paradox of the great speciators relies on knowledge of those genes in multiple systems. As discussed earlier, the candidate gene approach has many limitations and often shows conflicting results. Alternative hypothesis-free approaches, like genomewide association studies (GWAS), partially overcome the obstacles imposed by the incomplete understanding of the mechanisms underlying complex behaviours. GWAS requires phenotypes, in this case dispersal propensity, to be characterised at the individual level. Tasmanian migrants were relatively easy to identify because of their different morphological differences from the mainland subspecies. However, assigning a dispersal score to individuals where morphology cannot be used to distinguish between dispersers and non-dispersers becomes challenging. Dispersal phenotypes could be assessed via use of individual tracking devices though the sample sizes for these types of studies are often smaller than required for GWAS in particular. Genomics alone is unlikely to solve the complexity of the migratory phenotype. Recent studies have begun to incorporate transcriptome analysis in an effort to discover genetic determinants of migratory behaviour in birds, with the discovery of novel genes and chromosomal regions linked to migration strategies (Franchini et al., 2017; Lundberg et al., 2017; Horton et al., 2019; Frias-Soler et al., 2020). Regulatory regions with functional relevance have also been suggested to underlie many traits (Sackton et al., 2019). ATAC-seq offers the opportunity to explore genome-wide chromatin accessibility and transcription factor binding sites (Buenrostro et al., 2015).

Even though our results suggest that migrants have genetic differences from non-migrant individuals and that more dispersive populations have different *CREB1* profiles to non-dispersive ones, a more thorough sampling involving a higher number of loci is necessary to explore whether standing genetic variation within a population or *de novo* mutations can provide the raw material for natural selection to act upon shifting a population to complete sedentariness.

## Conclusions

In this study we assessed variation in six personality-related candidate genes in silvereye populations to examine whether signatures consistent with a genetic switch can explain rapid shifts in dispersal, leading to reduced gene flow and ultimately divergence in this great speciator. We find strong support for the idea that more dispersive populations carry longer *CREB1* alleles, but length decreases with time and limited isolation, suggesting that selection could be acting against dispersal ability following island colonisation. At the individual-level, partial migrants showed longer *CLOCK* alleles than non-partial migrant individuals. Our results suggest that the paradox of the great speciators can be partially understood from a genetic perspective.

## ACKNOWLEDGEMENTS

The samples used in this study were collected over two decades and we thank the many people who helped in numerous ways to facilitate the work. We are grateful to the chiefs and landholders of Vanuatu and New Caledonia for granting access to field sites, and field and logistic assistance from many people including the following: N. Clark, D. Treby, J. LeBreton, F. Cugny, W. Waheoneme, O. Hébert and A. Rouquié (New Caledonia); D. Treby, E. Sandvig and A. Robertson (Heron Island); S. Geiger, O. Boissier, R. Hills, S. Totterman, O. Drew, K. Ser, W. Ser, J. Saksak (Vanuatu), I.P.F. Owens, N. Clark (Australian mainland); J. Kikkawa (Norfolk Island); I.P.F. Owens (Lord Howe Island); P. Park, A. Fletcher, P. Gray (Tasmania); P. Schweigman, D. Onely (New Zealand); M. Bell (Chatham Island). A. Phillimore, R. Black, and N. Clark, kindly provided additional samples. The work was conducted under permits from the Direction de l’Environnement Province Sud and Direction Du Developpement Economique (New Caledonia and Loyalty Islands with thanks to G. Kakue); Vanuatu Environment Unit letters of permission and permits provided by E. Bani and we further thank D. Kalfatak and T. Tiwok for their assistance (Vanuatu); Lord Howe Island Board; Environment Australia (Norfolk Island); Queensland Department of Environment and Resource Management; Parks and Wildlife Service (Tasmania); NSW Office of Environment and Heritage (SL101536); New Zealand Department of Conservation Te Papa Atawhai; and Australian Bird and Bat Banding Scheme project and individual permits to SMC, MM, DP and BR. Ethics clearances were provided by University of Queensland ethics committee (ZOO/165/94/ARC, ZOO/520/96/ARC/PHD,ZOO/520/97/ARC/PHD) and Griffith University ethics committee (ENV/01/12/AEC, ENV/07/16/AEC, ENV/06/20/AEC, ENV/24/13/AEC) to SMC, and Charles Sturt University (15/024) to MM. We thank the funders of this work: Marsden Fund New Zealand to BCR and SMC to support fieldwork on mainland Australia and Heron Island and the molecular work; National Geographic Society Committee for Research and Exploration Grant (9383-13) to SMC to support fieldwork in New Caledonia; Natural Environment Research Coucil (NERC) postdoctoral fellowship to SMC to support fieldwork in Vanuatu and New Caledonia; AER was supported by a NERC studentship (NE/S007474/1) and a St John’s College Graduate Scholarship.

## Data and code availability

Code to reproduce all analyses is available at https://github.com/andreaestandia/1.0_silvereye_candidate_genes

