## Supplementary figures and images for "Candidate gene length polymorphisms are linked to dispersive behaviour: searching for a mechanism behind the “paradox of the great speciators”"

### Supplementary Figure 1

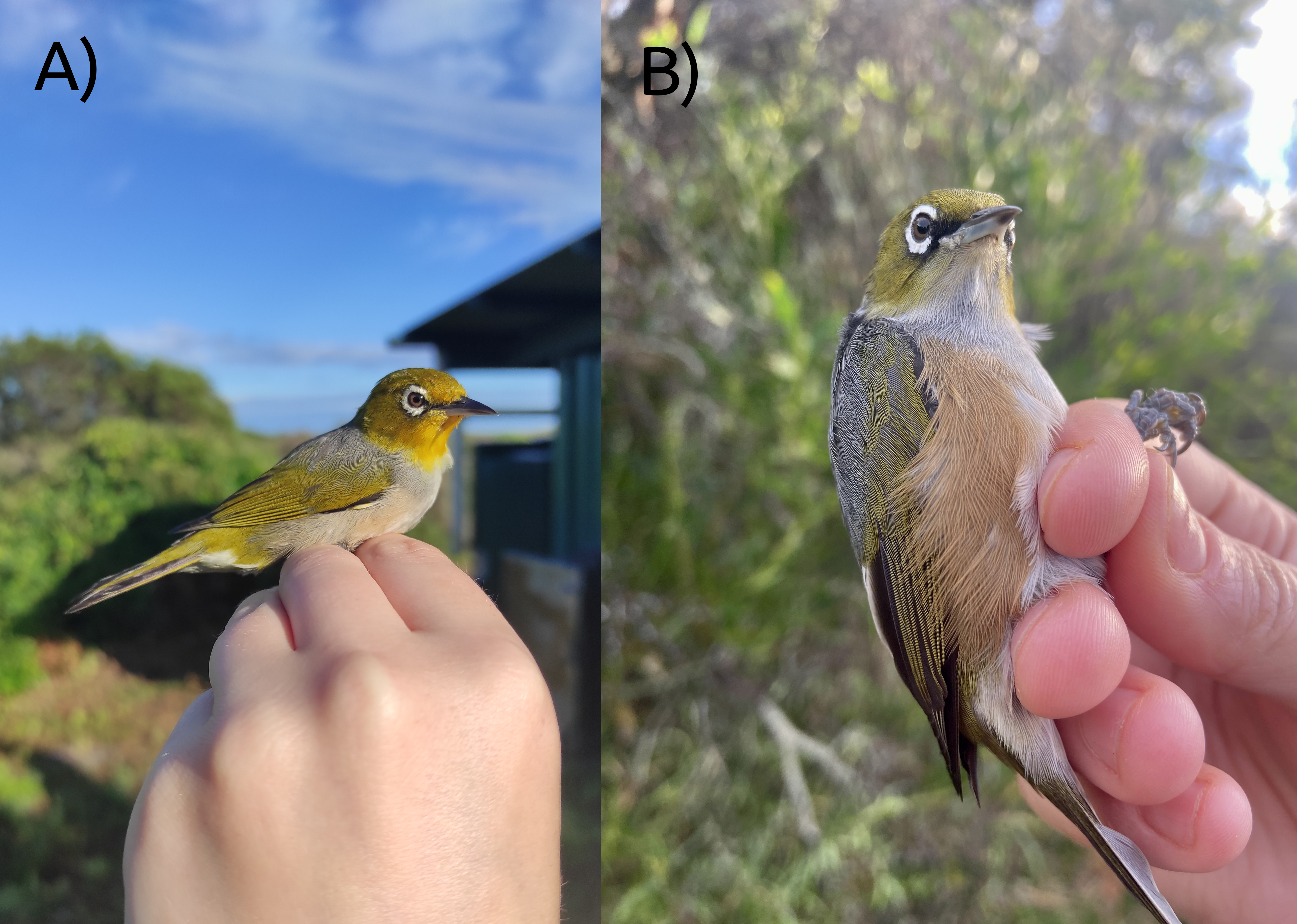

### Supplementary Figure 2

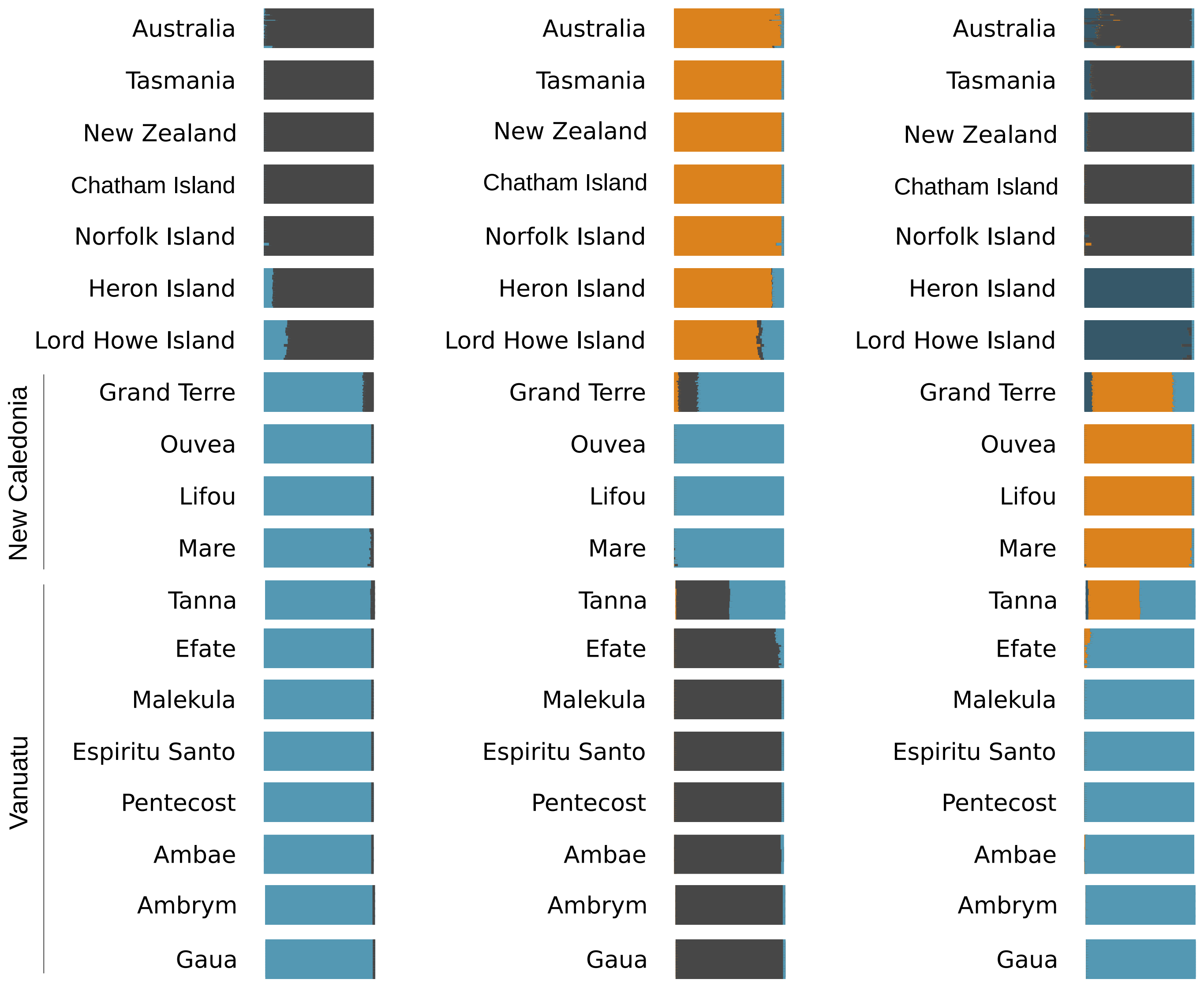

### Supplementary Figure 3

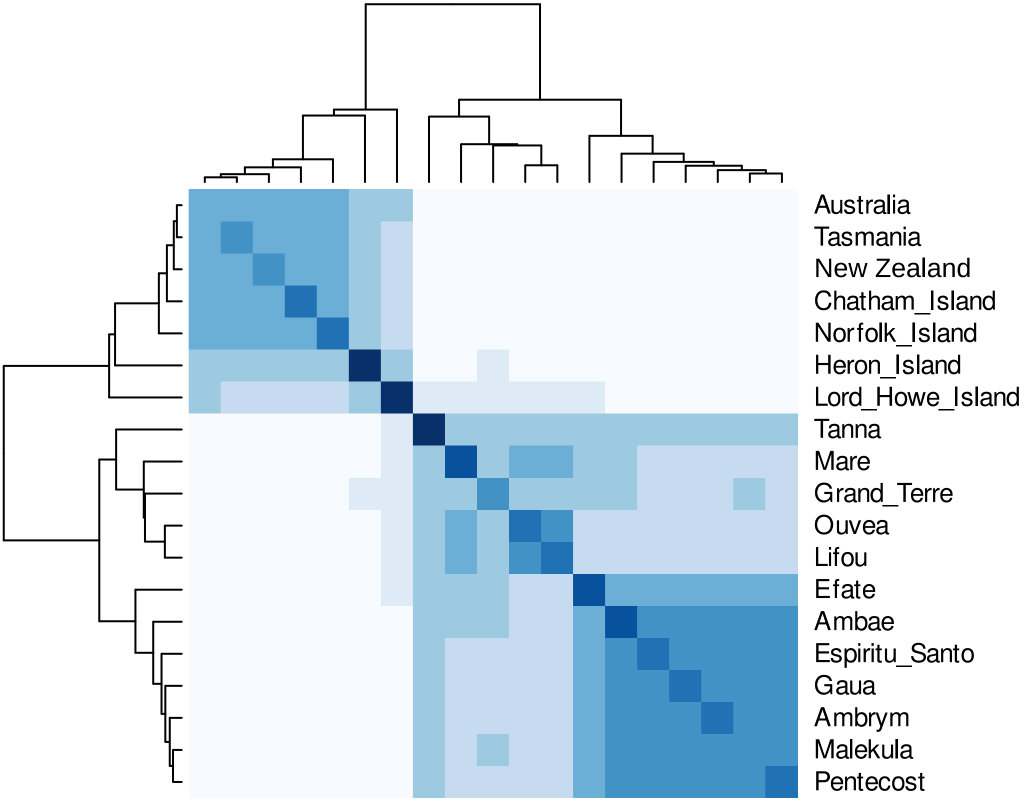

### Supplementary Figure 4

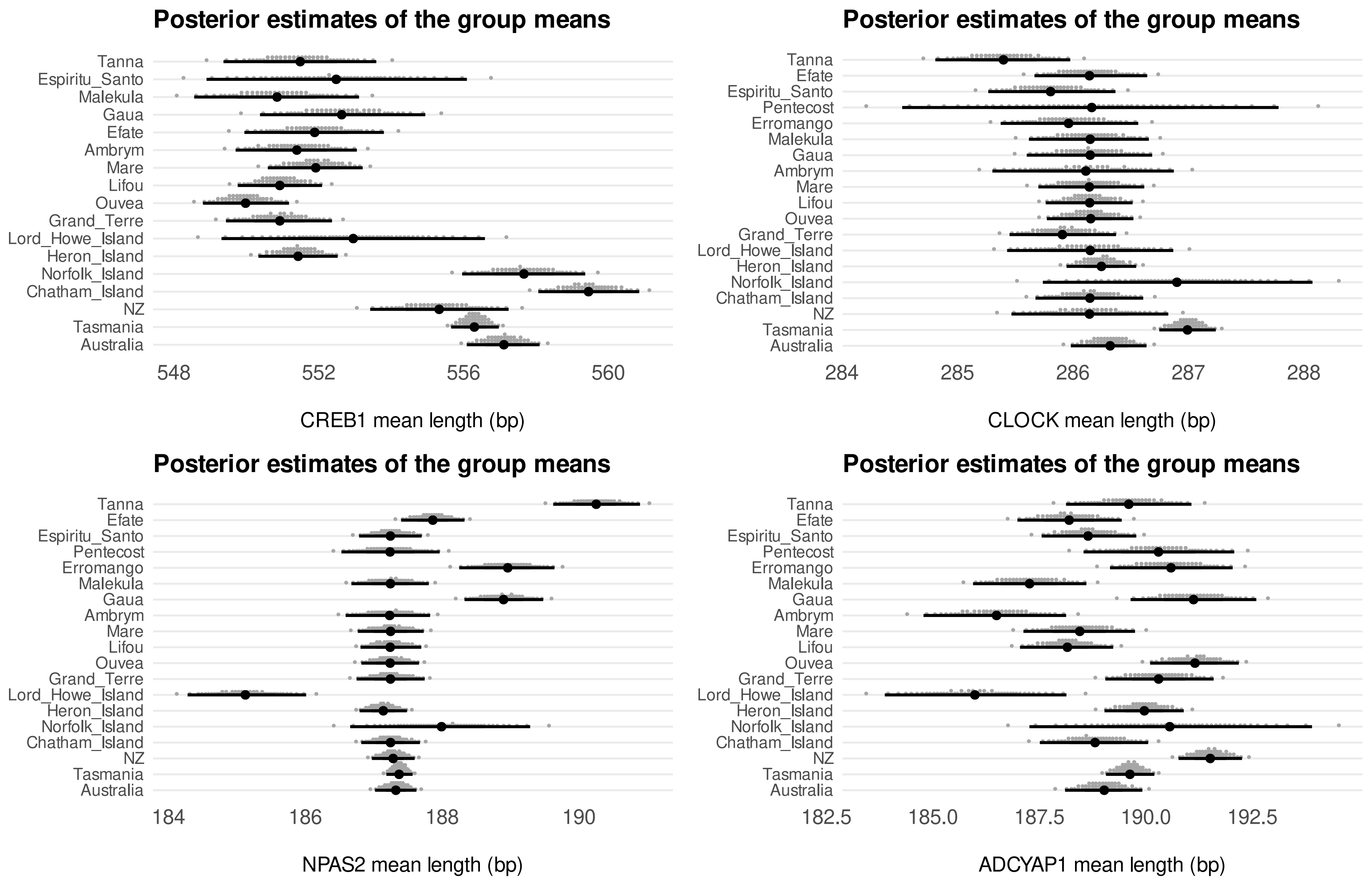
