## Supplementary Figures captions for "Candidate gene length polymorphisms are linked to dispersive behaviour: searching for a mechanism behind the “paradox of the great speciators”"

Supplementary figure captions

Figure S1. A) *Zosterops lateralis cornwalli* characterised by a bright solid yellow throat and grey flanks. This subspecies is resident on the east coast of the Australian mainland and coastal islands. Picture by Andrea Estandía on Broughton Island, New South Wales B) *Zosterops lateralis lateralis,* characterised by a grey throat and red flanks. This subspecies is found in Tasmania but some individuals migrate to the Australian mainland during winter. Picture by Andrea Estandía in Eagleby Wetlands, Queensland.

Figure S2. NGSadmix results of WGS data for *k=2* (left)*, k=3* (centre)*,* and *k=4* (right)*. k=2* represents the split between the ANZO and SM clusters, *k=3* the split within SM: New Caledonia and Vanuatu, although the southern island of Tanna, which is close to New Caledonia, has genetic affiliations with populations from both archipelagos.

Figure S3. Heatmap produced with the population-level covariance matrix generated with PCAngsd using WGS data. The divergence patterns are consistent with those from NGSadmix.

Figure S4. Posteriors were obtained extracted from the *brms* models. The point represents the median estimate and 89% credibility intervals are shown.
